# Removal of the catalytic subunit of DNA-protein kinase in the proximal tubules promotes DNA and tubular damage during kidney injury

**DOI:** 10.1101/2024.08.22.609216

**Authors:** M. Swanson, J. Yun, D.M. Collier, L. Challagundla, M. Dogan, C. Kuscu, M.R. Garrett, K.R. Regner, J. H. Chung, F Park

## Abstract

Tubular epithelial cell damage can be repaired through a series of complex signaling pathways. An early event in many forms of tubular damage is the observation of DNA damage, which can be repaired by specific pathways depending upon the type of genomic alteration.. In this study, we report that the catalytic subunit of DNA protein kinase (DNA-PKcs), a central DNA repair enzyme involved in sensing DNA damage and performing double stranded DNA break repair, plays an important role in the extent of tubular epithelial cell damage following exposure to injurious acute and chronic stimuli. Selective loss of DNA-PKcs in the proximal tubules led to increased markers of kidney dysfunction, DNA damage, and tubular epithelial cell injury in multiple models of acute kidney injury, specifically bilateral renal ischemia-reperfusion injury and single dose of cisplatin (15 mg/kg IP). In contrast, in a mouse model of kidney fibrosis and chronic kidney disease (UUO),the protective effects of DNA-PKcs was not as obvious histologically from the tissue sections. In the absence of proximal tubular DNA-PKcs, there was reduced levels of fibrotic markers, α-SMA and fibronectin, which suggests that there may be a biphasic role of DNA-PKcs depending upon the conditions exerted upon the kidney. In conclusion, this study demonstrates that the catalytic subunit of DNA-PKcs plays a context-dependent role in the kidney to reduce DNA damage during exposure to various types of acute, but not chronic forms of injurious stimuli.

## Introduction

The kidney is a major clearance organ in the body, which leads to the constant exposure of both endogenous and exogenous stimuli that can pose varying amounts of damage to the tubular epithelia. Injury can be acute (e.g. AKI) due to various factors, including infection, medication, and toxins, or chronic (e.g. CKD) due to hypertension or diabetes. In some cases, sub-lethal injury to the tubular epithelia can be self-limited, while in others, the severity is such that the kidneys are unable to fully recover. During these conditions, tubular epithelial cell remodeling may occur and initiate fibrotic pathway activation leading the kidney towards a progressive loss in organ function [1–3]. Although the cellular composition of the kidney is diverse and heterogeneous, the proximal tubular epithelia is a major site of investigation after injury due to their abundance and central role in the maintenance of kidney function. A common characteristic observed in different types of kidney injury is the detection of newly formed damage to the DNA [4–9]. Activation of DNA repair systems following oxygen or energy deprivation has been shown to have a positive influence on cell survival [6–8], but this may be pathway-dependent as not all DNA repair systems promotes a beneficial effect [10, 11]. DNA damage can induce various types of aberrant changes, including individual base alteration, pyrimidine dimer formation or single or double strand breaks. Of which, DNA double strand breaks are considered the greatest threat to genome stability that can determine the fate of the cell [12]. In the face of unrepaired DNA, it is likely that these cells undergo cell cycle arrest and depending upon the extent of the repair, the fate of the cell may require elimination through apoptotic cell death [13].

The catalytic subunit of DNA protein kinase (DNA-PKcs) is part of the phosphoinositide 3-kinase–like kinases (PI3KK) family, which also includes Ataxia telangiectasia and Rad3 related (ATR) and Ataxia telangiectasia mutated (ATM), and are serine/threonine kinases with distinct functions in the sensing and facilitating the activation of downstream effector systems to initiate the DNA repair process. In cultured kidney cells, cisplatin activated ATR to sense and initiate DNA repair [7]. A recent study in mice demonstrated that a selective ATR knockout in the proximal tubules prevented the tubular epithelial cell recovery following acute kidney injury (AKI) and accelerated fibrosis progression [5]. Unlike ATR, our knowledge about the role of DNA-PKcs in the kidney remains limited, particularly in the context of fibrotic progression. Genetically defective or absent DNA-PKcs causes severe combined immunodeficiency (SCID) due to its essential role in double stranded DNA break repair and immune cell diversity [2, 14]. In head and neck cancer, there is evidence that DNA-PKcs can promote DNA repair and accelerate cell proliferation through an interaction with thyroid receptor interacting protein 13 (TRIP13) [15]. Recent evidence in our lab has shown that reduced TRIP13 levels in the kidney during AKI can prolong tubular epithelial cell damage [16], whereas over-expression of TRIP13 in the kidney can correspondingly reduce kidney damage due to cisplatin exposure [4]. The potential interaction of TRIP3 with DNA-PKcs has not been investigated in the kidney during various types of injury.

In this study, a proximal tubule-selective knockout of DNA-PKcs was generated using the GGT1-Cre recombinase mouse to investigate how a segment-specific knockout of DNA-PKcs would affect the extent of DNA damage and tubular epithelial cell injury in various types of acute (cisplatin nephrotoxicity and bilateral ischemia-reperfusion injury) and chronic kidney injury (unilateral ureteral obstruction). In addition, the effect of biological factors, such as water restriction, diet and genetic interaction with TRIP13, was studied to determine whether they had any influence on DNA-PKcs function during kidney injury. Cumulatively, our findings suggest that after tubular injury, DNA-PKcs plays a central role in reducing DNA damage to lessen the renal tubular epithelial cell injury during exposure to various types of AKI insults, but its role may not be as relevant during the progressive phases of kidney fibrosis.

## Materials and Methods

### Generation of proximal tubule DNAcs knockout and Trip13 over-expression mice

Floxed DNA-PKcs^fl/fl^ mice were previously published [17] and generously provided by Dr. Jay H. Chung (NIH). Conditional proximal tubule-expressing GGT1-Cre^+/-^;*Trip13*^Stop/Stop^ (*Trip13*^ΔStop^) transgenic mouse were generated using a floxed stop strategy (Cyagen, Santa Clara, CA) as recently published by our lab [4]. Representative PCR genotyping from genomic DNA isolated from the litters prior to experimental use are shown in Supplemental Figure for DNA-PKcs. Male and female DNA-PKcs^fl/fl^ and GGT1-Cre^+/-^;*Trip13*^Stop/Stop^ mice were interbred to produce double GGT1-Cre^+/-^;DNA-PKcs^fl/+^ or triple heterozygote GGT1-Cre^+/-^;DNA-PKcs^fl/+^;*Trip13*^Stop/+^ mice. Further interbreeding with GGT1-Cre^-/-^;DNA-PKcs^fl/fl^, or GGT1-Cre^-/-^;DNA-PKcs^fl/fl^;*Trip13*^Stop/Stop^ was performed to produce the experimental mice with the appropriate genotypes: GGT1-Cre^+/-^;DNA-PKcs^fl/fl^ (DNA-PKcs^PTKO^) and GGT-Cre^+/-^;DNA-PKcs^fl/fl^;*Trip13*^Stop/Stop^ (DNA-PKcs^PTKO^; *Trip13*^ΔStop^). For genetic controls denoted as DNA-PKcs^CTRL^, the following genotypes were combined, GGT1-Cre^+/-^;DNA-PKcs^+/+^ and GGT1-Cre^-/-^;DNA-PKcs^fl/fl^, as no obvious difference was noted in their phenotyping. Mice were housed, bred and maintained in a specific pathogen-free animal facility at the University of Tennessee Health Science Center in Memphis. All experiments were approved by the Institutional Animal Care and Use Committee (IACUC #19-0102 and #21-0223). Mice were allowed *ad libitum* access to food and water, and maintained in a 12 hr light/12 hr dark cycle. Genotypes were confirmed by PCR amplification of genomic DNA isolated from toe or tail snip as previously described by our lab [4] or the Chung lab [17].

### Murine model of cisplatin-induced nephrotoxic AKI

Male DNA-PKcs^PTKO^ and DNA-PKcs^PTKO^;Trip13^ΔStop^ and genetic control DNA-PKcs^CTRL^ mice (aged 8–12 weeks) were injected with a single intraperitoneal dose of cisplatin (15 mg/kg) (Advanced ChemBlocks, cat #K12017) to produce cisplatin nephrotoxic AKI. As a control, vehicle (sterile saline) was injected with the same range of volume used in the cisplatin solution. To determine whether volume status or dietary fat intake affected the DNA damage induced by cisplatin, mice were water restricted for different lengths of time prior to the injection of cisplatin: 1) 7-8 hours or 2) 12-16 hours. In some mice, we provided high fat (HF) diet (60% kcal, cat #D12492, Research diets) to compare with their normal fat (NF) diet (Inotiv 7912, 17% kcal). Body weights were measured before and after water restriction, and every 24 hours following injection and continued to the end of the experimental period.

### Murine model of bilateral ischemia-reperfusion injury (IRI)

Male DNA-PKcs^PTKO^ and PKcs^PTKO^; *Trip13*^ΔStop^ and genetic control DNA-PKcs^CTRL^ mice (aged 8–12 weeks) were randomly distributed in either sham or surgical AKI operated groups. Body weights were measured prior to two separate injections with an anesthesia mixture containing ketamine (100 mg/kg IP) and xylazine (20 mg/kg IP), and a pain reliever containing slow release buprenorphine (0.1 mg/kg IP). After the mice were fully under anesthesia, the mice were prepared for sterile surgery. A flank incision was made on both the left and right side of the body to expose the kidneys for the isolation of the renal artery and vein and the separation of the renal nerves. A clamp was placed onto the isolated renal vessels for 21.5 minutes and the kidneys were placed back into the abdominal space for the duration of the ischemic period. Afterwards, the clamps were removed and the wounds were closed. Thereafter, the mice were allowed recover in their home cage for the 24 hour experimental period.

### Murine model of unilateral ureteral obstruction (UUO)

Male and female PKcs^PTKO^ and genetic control DNA-PKcs^CTRL^ mice (aged 8–12 weeks) mice were randomly distributed (aged 8–12 weeks) in either sham or UUO-operation groups, and DNA-PKcs^PTKO^;Trip13^ΔStop^ was used for UUO surgeries. Body weights were measured prior to two separate injections with an anesthesia mixture containing ketamine (100 mg/kg IP) and xylazine (20 mg/kg IP), and a pain reliever containing slow release buprenorphine (0.1 mg/kg IP). After the mice were fully under anesthesia, the mice were prepared for sterile surgery. A flank incision was made in the left side of the body to expose the kidney and isolate the ureter to place two distinct ligatures using 5/0 silk to stop urine flow. The kidney was placed back into the abdominal space, the wounds were closed, and the mice recovered in their home cage for the duration of the experimental period of either 7 or 14 days.

### Sample collection and analysis

After the surgical procedures, body weights were measured every 24 hours for the first 7 days, and afterwards every other day for the 14 day period. Blood was collected prior to the collection of the kidneys and the serum was isolated for measurement of creatinine by LC/MS/MS (Department of Biochemistry, University of Alabama-Birmingham). The kidneys were removed and placed in either neutral buffered formalin for paraffin embedding to perform immunohistochemistry and tissue staining, or on dry ice for molecular and protein analysis.

### Histopathology

Paraffin embedded sections (4 µm) were deparaffinized in xylene and rehydrated in increasing ethanol percentages to prepare for staining with hematoxylin and eosin (H&E). The kidney sections were de-identified and scored by a blinded nephrologist with expertise in rodent models of AKI to assess tubular injury. The criteria for damaged tubules included the identification of flattened epithelia, tubular dilation and cast formation as previously described in our lab [15, 16, 18–21]. The injured tubules were calculated as a percent of the total number of tubules counted in 5 different sections from every animal in each group.

### Antibodies

Rabbit anti-phospho-H2A.X (Ser139) (Cell Signaling cat #9718), phospho-Histone H3 (Ser10; cat #9701; 1:250; Cell Signaling), cleaved caspase-3 (cat #9661, 1:200; Cell Signaling), and Ki67 (cat #15580, 1:200; Abcam), rabbit anti-phospho- (Cell Signaling cat #9101) and total ERK1/2 (Cell Signaling cat #9102), rabbit anti-TRIP13 (Abcam cat #128153), rabbit anti-MLKL (Cell Signaling cat #37705) and phospho-MLKL (Ser345) (Cell Signaling cat #62233), and mouse anti-α-smooth muscle actin (1A4) AlexaFluor 488 (Clone 1A4; Invitrogen cat #53-9760-82) were purchased. As a loading control, β-actin or GAPDH (1:1,500; Cell Signaling) was used. Secondary goat anti-rabbit IgG (Invitrogen cat # G21234) and goat anti-mouse IgG conjugated to horseradish peroxidase (HRP) (Cell Signaling cat #7076). All antibodies have been previously used in published studies demonstrating that other labs have confirmed their specific target.

### Immunofluorescent histochemistry in kidney sections

Immunohistochemistry was performed on deparaffinized kidney sections (4 µm). The kidney sections were deparaffinized in xylene and rehydrated through a series of decreasing percentages of ethanol (100% to 70%) Antigen retrieval was performed by heating the slides for 45-50 minutes in epitope retrieval solution (cat # IW-1100; IHC World) slides were washed using 1X TBS between each of the steps listed below (cat# 28358; Thermo Scientific). Slides were then incubated in peroxidase blocking solution for 15 minutes. The tissue sections were washed in 1X TBS with 0.08% Triton X for 10 minutes, and blocked in 2.5% normal donkey serum for 60 minutes. Polyclonal Ki67 (cat #15580, 1:200; Abcam) were diluted in 1X TBS with 0.08% Triton X and 2% donkey serum (cat# 50062Z; Life Technologies) and was incubated on the tissues overnight at 4°C. The tissues were washed 3x in 1X TBS, and the secondary donkey anti-mouse IgG conjugated with AlexaFluor 555 (1:250 dilution; cat #A-31570; Invitrogen). To determine co-localization with the proximal tubule, biotinylated *Phaseolus vulgaris* hemoagglutinin-E (PVA-E; 1:500 dilution; proximal tubules; cat # B-1125; Vector Laboratories) or tubulointerstitial cells, α-SMA (1:500), was incubated onto the sections for 30 minutes at 25°C, and then with streptavidin-linked IgG with AlexaFluor 488 (1:500 dilution, cat # S11223; Invitrogen) for 30 minutes. The sections were mounted with DAPI (cat # H-1200; Vector Laboratories) and imaged on the EVOS light microscope at 20-40x magnification.

### Immunohistochemistry using DAB as the substrate

Immunohistochemistry was performed [16, 20, 22] on deparaffinized kidney sections (4 µm). In brief, antigen retrieval was performed by heating for 50-60 minutes in epitope retrieval solution (cat# IW-1100; IHC WORLD). As the previous section stated the slides were washed with 1X TBS between each of the steps below. Slides were then incubated in peroxidase blocking solution to minimize endogenous peroxidase activity, and the tissue was then permeabilized in 1X Tris buffer saline containing 0.08% Triton-X (cat# T8787; Sigma). The following primary polyclonal rabbit antibody to phospho-H2AX (Ser139) (1:250), phospho-Histone H3 (Ser10) (1:250), cleaved caspase-3 (1:200), and Ki67 (1:200) were used on the kidney sections. Secondary goat anti-rabbit IgG (1:250) conjugated to horseradish peroxidase (HRP) (1:250) were used and detected using diaminobenzidine (DAB; cat # D8001; Sigma, St. Louis, MO). Sections were counterstained with hematoxylin prior to the cover slipping protocol. Images were taken with an EVOS light microscope at 20-40 fold magnification. The percentage of positive nuclei for the target proteins were calculated as a total number of nuclei.

### Western blot analysis

Protein lysates were isolated from the harvested mouse kidneys by homogenization in 1X RIPA buffer (ThermoFisher) containing protease and phosphatase inhibitors (Pierce) followed by differential centrifugation as previously described in our lab [4, 16, 20, 22, 23]. Western blot analysis was performed using a gradient SDS-PAGE gel (4-12%), transferred to either PVDF or nitrocellulose membranes, and bands were detected by chemiluminescence (GE) using Bio-Rad ChemiDoc MP imaging system. All of the antibodies were tested using proteins transferred onto PVDF membranes except for cyclin B1 (nitrocellulose).

### Sirius red staining

Sirius red staining was performed on the kidney sections using a modified protocol from Polysciences. In brief, the kidney sections were deparaffinized and dehydrated through a series of decreasing percentages of ethanol (100% to 70%), washed in distilled water, and then stained with Sirius red solution for 60 minutes. The slides were immediately washed in water, placed in 75% ETOH for 45 seconds, then rehydrated with increasing concentrations of ETOH and xylene. All of the slides were coverslipped and imaged using a light microscope for analysis of collagen deposition. Red staining was analyzed as previously described in our lab [24].

### Single nuclei RNASeq extraction, library construction, and DNA sequencing

Initially, mouse kidney tissues are cut into small pieces to extract nuclei using Nuclei EZ Lysis buffer (NUC-101; Sigma-Aldrich) containing protease (Sigma-5892791001) and RNase inhibitor (N2615, Promega; AM2696, Life Technologies) with 10 times of pestle A and B treatment for each. Digested tissues were incubated in this buffer on ice for 5 minutes. Post incubation, the samples were passed through a 40 µm cell strainer (43-50040-51; pluriSelect) and subjected to centrifugation at 500 × g for 5 minutes at a temperature of 4°C. Subsequently, the sediment was mixed again with 2mL of the same buffer and processes with pestle B 10 times. Nuclei were incubated once more on ice for 5 minutes. A repeat centrifugation was carried out, after which the sediment was mixed in Nuclei Suspension Buffer (PN-2000153; 10x Genomics; containing 1x PBS, 1% bovine serum albumin, 0.1% RNase inhibitor) and passed through a filter as per the initial method. The segregated nuclei were then stained with DAPI and quantified using the Countess 3 Automated Cell Counter (ThermoFisher). Finally, viable nuclei were diluted to 1000 cells/ul and combined with a barcode-labelled gel bead, utilizing the 10x Chromium system (10x Genomics). Total 7,000 nuclei were loaded for each mouse kidney tissue.

### DNA library preparation

Within each oil-droplet (GEM), RNA was reverse-transcribed to cDNA using the 10x Chromium Single Cell 3′ Library & Gel Bead Kit v3.1 (10x Genomics) as per the protocol provided by the company. After GEM-RT cleanup, cDNA amplified for 13 cycles due to generation of cDNA from nuclei. The cDNA’s concentration was gauged using a Qubit fluorometer (ThermoFisher), and approximately 100 ng (or 10-15 µl) was utilized for downstream process where cDNA underwent fragmentation, end-repair and A-tailing. This was followed by the affixing of Illumina adapters, in line with the procedures provided by the 10x Chromium Single Cell 3′ Library & Gel Bead Kit v3.1 (10x Genomics) by using SI-TT-double index for sample indexing. The reaction conditions were set at 98°C for 45 seconds, with 13 cycles at 98°C for 20 seconds, 54°C for 30 seconds, and 72°C for 20 seconds, concluding with a final extension at 72°C for 1 minute and a hold at 4°C. After the completion of the PCR step, the quantity of cDNA was appraised using a Qubit fluorometer (ThermoFisher). The expected size distribution post adapter ligation was confirmed using Agilent TapeStation. Multiplexed libraries were pooled by equal molarity for sequencing using 2X P3 (100 cycle kit) on NextSeq 2000 (Read 1:28 cycles, i7 index: 10 cycles, i5 index: 10 cycles, Read 2: 90 cycles) at the University of Mississippi Medical Center-Molecular and Genomics Core Facility.

### Sequence Alignment and Analysis

The raw sequencing reads were processed using Cell Ranger v7.0 (10x Genomics). The Cell Ranger mkfastq pipeline was used to demultiplex the FASTQ base call files. The outputs were then used to run Cell Ranger count which align the sequencing reads to the custom mouse reference transcriptome to include the transcripts for GFP and Cre. The outputs from Cell Ranger for all samples were merged together into R(v4.3.0) [25] environment using Seurat (v5) [26]. The next step involved quality filtering including but not limited to finding cells which are outliers with respect to the rest of the dataset such as low-quality cells, doublets/multiplets (DoubletFinder v2.0.4) [27], high mitochondrial gene content etc. followed by the FindVariableFeatures() function with defaults which applies variance-stabilizing transformation. The data was normalized using the sctransform () flavor V2 approach as described in Choudhary and Satija ([28]) as is needed before performing Principal Component (PC) Analysis. By default, Seurat only uses the variable features determined in the previous step as input. Using the new streamlined integration function (IntegrateLayers() and specifically integrating using the method “Harmony”) we obtain a Seurat object with the original count matrix as well as an “integrated” matrix with batch corrected counts. Using the integrated and filtered matrix, FindNeighbors() was run which uses the first 30 principal components (default) to construct a cellular KNN graph based on the Euclidean distance in PCA space and then edge weights between the cells are refined based on Jaccard similarity. The cells were then iteratively clustered into clusters by applying the Louvain algorithm. The resolution parameter determines the granularity expected when grouping cells together (we used 0.2 for the preliminary results). The same PCs as calculated before were used as input for generating a two-dimensional visualization of the cells grouped by cluster using Uniform Manifold Approximation Projection (UMAP). The marker genes for each cluster were calculated through the FindAllMarkers() function to calculate the overexpressed genes in each cellular cluster compared to all other clusters using the canonical markers for kidney cell types. The violin plots for these genes showing their expression across the cell types were generated. Differential gene expression analysis was conducted between all pairs of groups among the proximal tubule as well as the injured proximal tubule cell type.

### Cell counting assay

Mouse TKPTS cells (ATCC cat #CRL-3361) in DMEM/F-12 (cat# 11300-032; GIBCO) were plated into 6-well dishes and counted by hemocytometry to provide the initial time point of 0 hour values. The cells were treated with either vehicle or drugs for 24 hours at which point the cells were counted at 24 and 48 hours by hemocytometry. The cell counting was performed with n=2-3 samples/experiment and replicated on 4 different attempts, and the average values from each experiment were compiled and averaged to determine the overall changes in cell numbers.

### Statistics

All data was shown as mean ± standard error mean (SEM). The data were tested for normality with the Kolmogorov-Smirnov test using Prism 6.0. Normal distributed data was tested with analysis of variance (ANOVA) and subsequent post-hoc analysis was performed by Tukey’s multiple comparison test. Non-normal distributed data were tested with the Kruskal-Wallis test for significance among multiple groups followed by Dunn’s multiple comparison test. P value ≤0.05 was considered significant. For the differential gene expression analysis, all pairs of groups among the proximal tubule as well as the injured proximal tubule cell type were compared. Pathway enrichment analysis was performed in Reactome [29] for differentially expressed genes using a log2FC cutoff at 0.5.

### Study approval

All animal protocols (AUA #17-032, #18-097, #19-0102, and #21-0102) were approved by the UTHSC IACUC prior to performing any of the experiments.

## Results

### Role of DNA-PKcs on kidney injury in a mouse model of cisplatin-induced AKI

In a prior study in our lab, we found that there was a significant increase in DNA damage, tubular damage and reduced renal function using NU-7441 (10 mg/kg IP), which is a highly selective inhibitor to DNA-PKcs, in cisplatin nephrotoxicity [4]. In that study, the solubility of NU-7441 was very poor, so the formulated solution required a relatively high volume (∼0.8-1 mL) to administer the full dose into each mouse and led to only a modest difference in the extent of kidney damage between the groups treated with cisplatin along with the vehicle solution. Regardless, pharmacological inhibition of DNA-PKcs was capable of significantly increasing DNA damage by ∼14-fold in the cisplatin-treated mice co-administered with NU7441 compared to those mice treated with cisplatin and vehicle solution. For this reason, we postulated that the volume status at the time of cisplatin administration would be an important factor to consider in the experimental design for the current study and so we examined shorter and longer periods of water restriction prior to the administration of cisplatin.

Regardless of whether the DNA-PK^CTRL^ mice were withheld water for either short (7-8 hours) or long (12-16 hours) period prior to an injection of cisplatin, whole body weight tended to be decreased over the 72 hours experimental period and used as a proxy for their general health (**Fig. 1A-1C**). Vehicle-treated mice after water restriction regained their pre-thirsted weight and did not exhibit any loss in body weight. Weight loss of the mice did not necessarily correlate with reduced kidney function as measured by serum creatinine levels (**Fig. 1D-1F**). No apparent effects on renal architecture was observed between the short or long water restricted mouse kidneys (**Fig. 1G-1P**). The only cisplatin-treated group that did not Serum creatinine levels were not significantly different between vehicle- or cisplatin-treated mouse groups fed NF diet and thirsted for only 7-8 hours (**Fig. 1D** and **1E**). However, serum creatinine was elevated if NF-fed mice were water restricted for a longer period prior to cisplatin injury (**Fig. 1F**) or if the cisplatin-treated mice were fed a HF diet (**Fig. 1D**) under the shorter water restriction condition.

**Figure 1.** Increased DNA damage and kidney injury in proximal tubule-specific knockout of DNA-PKcs in mice. Mice with various DNA-PKcs^CTRL^ or DNA-PKcs^PTKO^ genotypes were placed on a normal fat (NF) and high fat (HF) diet at 4 weeks of age for a period of 6 weeks. Prior to the end of the experimental period, mice were injected with a single dose of cisplatin (C; 15 mg/kg IP) or vehicle (V; saline) and body weights were measured for the subsequent 3 days. These values are shown in the graphs for **Fig. 1A-1C**. At the end of the experimental period, serum creatinine was measured (**Fig. 1D-1F**) and kidneys were harvested to determine tubular damage by H&E staining. (**Fig. 1G-1P**). Representative kidney sections were shown from the various mouse kidneys. Short (7-8 hr) and long (12-16 hr) dehydration is noted in the graph and figures for the length of time prior to administering either vehicle or cisplatin prior to time 0. * P<0.05, ** P<0.01, *** P<0.001 indicates significant differences between groups. Control mice with the genotypes (DNA-PKcs^CTRL^) or proximal tubule-specific knockout (DNA-PKcs^PTKO^) and ρStop (DNA-PKcs^PTKO^;Trip13^ΔStop^). V-NF = vehicle normal fat diet; V-HF = vehicle high fat diet; C-NF = cisplatin normal fat diet; C-HF = cisplatin high fat diet. Scale bar = 200 µm.

In the mouse groups with the prolonged water restriction prior to cisplatin treatment (**Fig. 1F**), vehicle-treated DNA-PK^CTRL^ or DNA-PK^PTKO^;Trip13^ΔStop^ serum creatinine was measured at a low basal level (0.06 ± 0.01 mg/dL; n=6 and 0.05 ± 0.01 mg/dL; n=4, respectively) at day 3 after injection (**Fig. 1F**), and no loss in body weight over the 72 hour experimental period (**Fig. 1C**). Cisplatin treatment (15 mg/kg IP) to DNA-PK^CTRL^ mice led to a significantly >3-fold higher serum creatinine levels to 0.19 ± 0.05 mg/dL (n=9). In mouse kidneys with a combined loss of proximal tubular DNA-PKcs and increased expression of TRIP13 (DNA-PK^PTKO^;Trip13^ΔStop^), the kidney injury was partially mitigated as the creatinine levels were mildly, but not significantly increased to 0.09 ± 0.02 mg/dL (n=7) compared to the vehicle-treated mice at day 3. The creatinine levels were similar in vehicle- and cisplatin-treated DNA-PK^PTKO^;Trip13^ΔStop^ mice at day 1 (unpublished observations) with minimal DNA damage (0.15 ± 0.05%; n=4).

### Increased DNA damage in the kidney following cisplatin nephrotoxicity due to a high fat diet

To determine whether high fat could modulate the extent of DNA damage and kidney injury following cisplatin administration, we modified the dietary intake ingested by the mice after week 4 following birth. In the majority of the mice studied in our experiments, the normal fat (NF) diet typically provided by our animal facility was used during our protocol. To compare the fat content on cisplatin-dependent kidney effects, we used a high fat (HF) starting at week 4 after birth for a period of 6 weeks. In the final week of the experimental period, the mice were transiently withheld water (either 7-8 or 12-16 hours) prior to the administration of a single dose of cisplatin (15 mg/kg IP) or vehicle (saline). These mice were closely monitored over the next 72 hours and body weights were measured prior to the collection of blood and kidneys for analysis. In some mice, food intake and urine were collected using metabolic cages.

As shown in **Fig. 2**, DNA damage in mouse kidneys due to cisplatin administration appeared to be dependent upon the length of water restriction and the fat content in the mouse chow. Regardless of the length of water restriction prior to injecting vehicle (saline) solution, the basal percentage of γ-H2AX positive nuclei was very low in each of the groups regardless of the presence or absence of DNA-PKcs. In the 7-8 h water restricted mice fed NF (**Fig. 2A, B**) or HF (**Fig. 2C, D**), the percentage of γ-H2AX positive nuclei in DNA-PKcs^CTRL^ kidney sections was 0.3 ± 0.1% (n=10) or 0.2 ± 0.1% (n=5), respectively (**Fig. 2G**). In the absence of the proximal tubular expression of DNA-PKcs (vehicle-treated DNA-PKcs^PTKO^), mice fed fedNF or HF showed 0.9 ± 0.3% (n=4; **Fig. 2E** and **2H**) or 0.2 ± 0.03% (n=3; **Fig. 2H**), respectively, in γ-H2AX positive nuclei. In the longer 12-16 h water restriction group, vehicle injection did not affect the basal γ-H2AX positive nuclei (0.3 ± 0.1%; n=6; **Fig. 2I**) in DNA-PKcs^CTRL^ kidney sections compared to the shorter water restriction group.

**Figure 2.** Increased DNA damage due to cisplatin nephrotoxicity due to long-term high fat diet or loss of DNA-PKcs in the proximal tubules. 4 week old mice were either kept on normal fat diet (A, B, E, and F) or placed on a high fat diet (C, D) for an additional 6 weeks. (A-F) Representative kidney sections stained for γ-H2AX from DNA-PKcs^CTRL^ mice on a normal (A, B) or high fat (C, D) diet treated with vehicle (saline) (A, C) or cisplatin (15 mg/kg IP) (B, D). In (E, F), kidneys from DNA-PKcs^PTKO^ mice on normal fat diet treated with vehicle or cisplatin (15 mg/kg IP). (G-I) γ-H2AX positive nuclei are calculated as a percentage of total nuclei. Individual animals are shown in each plot. *P<0.05, ** P<0.01, **** P<0.0001 indicates significant differences between groups. Control mice with the genotypes (DNA-PKcs^CTRL^) or proximal tubule-specific knockout (DNA-PKcs^PTKO^) with DNA-PKcs^PTKO^;Trip13^ΔStop^over-expressing Trip13. NF = normal fat; HF = high fat; V = vehicle; C = cisplatin. Scale bar = 200 µm.

Following the single injection of cisplatin (15 mg/kg IP) after the mice were water restricted for 7-8 hours, there was a non-significant change to 1.1 ± 0.6% (n=4) in γ-H2AX positive nuclei for NF DNA-PKcs^CTRL^ mice kidneys (**Fig. 2B** and **2G**). In HF DNA-PKcs^CTRL^ mice, there was a modest, but significantly higher (P<0.0001) percentage of γ-H2AX positive nuclei (3.6 ± 0.5%; n=6) (**Fig. 2D** and **2G**), which correlated with a significantly higher serum creatinine compared to their corresponding HF control group (**Fig. 1D**). In the DNA-PKcs^PTKO^ kidney sections from NF-fed mice, cisplatin-treatment led to a considerably higher number of γ-H2AX positive nuclei (17.9 ± 3.3%; n=5; **Fig. 2F** and **2H**).

As mentioned in the Introduction, prior work by Banerjee et al. [15] in head and neck cancer cells demonstrated that TRIP13 promoted DNA repair through NHEJ by direct binding to DNA-PKcs. In our lab, we investigated the effect of a proximal tubule-specific TRIP13 over-expression (Trip13^ΔStop^) [4] by breeding into the DNA-PKcs^fl/fl^ mouse to determine the effect on DNA damage during UUO and its subsequent role, if any, in tubular remodeling and fibrosis progression. After 12-16 hours of water restriction prior to cisplatin injection, the γ-H2AX positive nuclei was significantly higher (P<0.001; 28.7 ± 3.2%; n=9) (**Fig. 2I**) compared to vehicle-treated DNA-PKcs^CTRL^ mice (0.15 ± 0.05% n=4). In the presence of DNA-PK^PTKO^;Trip13^ΔStop^ mouse kidneys where the DNA-PKcs is deleted with TRIP13 overexpression in the proximal tubules, there was a markedly reduced DNA damage by ∼70% down to 9.0 ± 1.9% (P<0.001; n=7) γ-H2AX positive nuclei. These finding suggest that the length of water restriction prior to administering the cisplatin can influence the extent of DNA damage in the kidney after 3 days, and TRIP13-overexpression even in the absence of DNA-PKcs can reduce DNA damage in the kidney.

### Epithelial cell proliferation and mitotic index

The tubular epithelial cell proliferation status as determined by Ki-67 positive (denoted as Ki67+) percentage was low in vehicle-treated DNA-PKcs^CTRL^ kidneys in both NF (1.8 ± 0.3%; n=10) and HF (1.2 ± 0.3%; n=6) groups (**Fig. 3A**). Genetic removal of DNA-PKcs in the proximal tubules (DNA-PKcs^PTKO^) did not have any effect on basal proliferation as theKi67+ nuclei were similarly low in HF (0.9 ± 0.3%; n=3) and NF (0.8 ± 0.1%; n=4) kidney sections (**Fig. 3D**). Cisplatin treatment did not markedly affect the Ki67+ cells in the NF-fed DNA-PKcs^CTRL^ mouse kidneys, but there was a mild, but significantly lower percentage of Ki67+ nuclei (P<0.05; n=6) in the kidneys from HF- versus NF-fed DNA-PKcs^CTRL^ mice. On the other hand, a modest increase in Ki67+ nuclei was detected following cisplatin treatment in the DNA-PKcs^PTKO^ kidneys (1.9 ± 0.8%; n=5), but this was not significantly different when comparing all kidney groups together (**Fig. 3A** and **3D)**.

**Figure 3.**
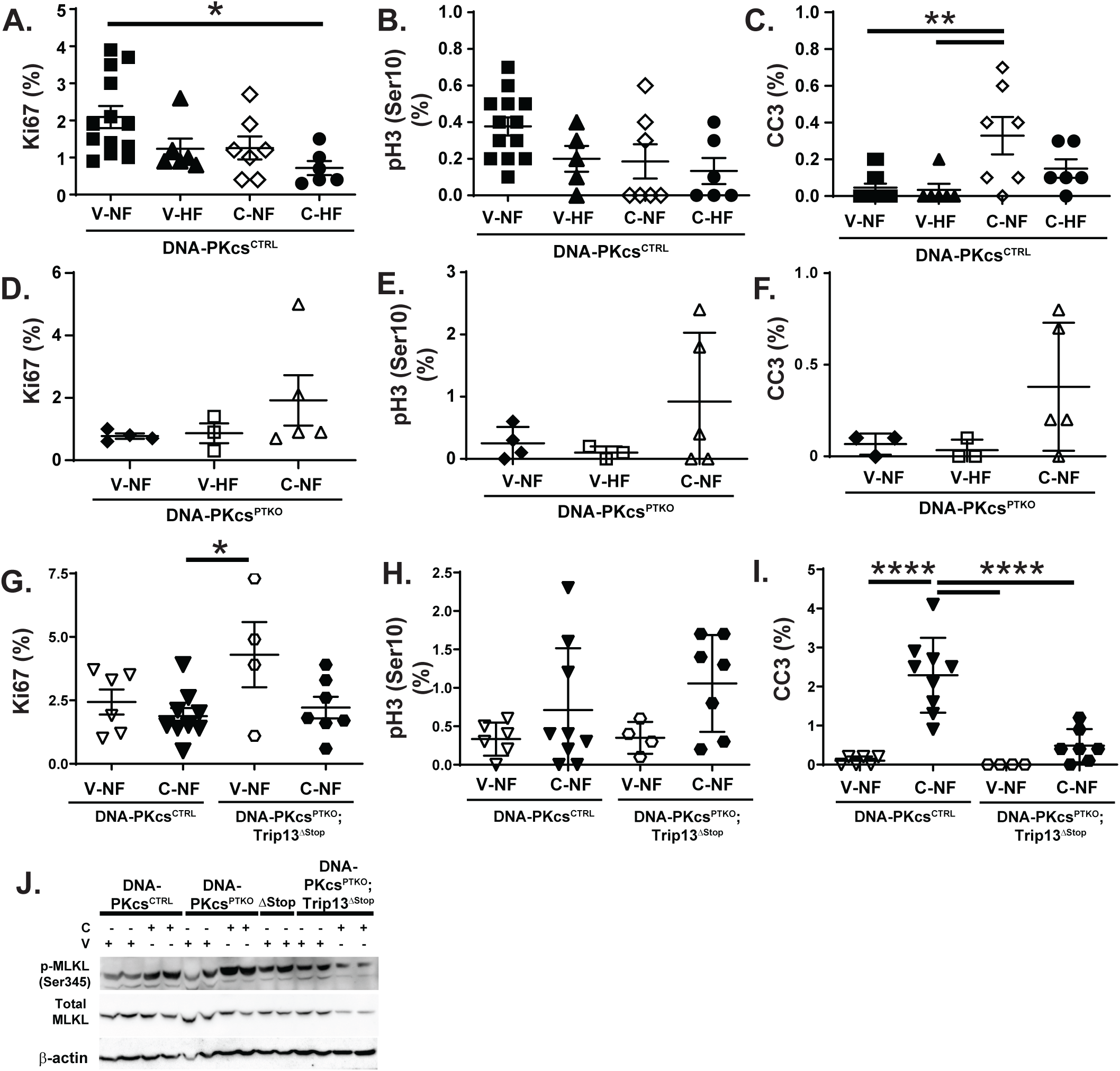
Effect of high fat or loss of DNA-PKcs on cell proliferation and programmed cell death in the kidney following cisplatin treatment. (**A, D, G**) Ki-67, (**B, E** and **H**) p-H3 (Ser10), and (**C, F,** and **I**) cleaved caspase-3 (CC3) positive nuclei were counted in kidney sections obtained from normal (NF) or high fat (HF) diet mice treated with a single injection of vehicle (saline) or cisplatin (15 mg/kg IP). (A-C) mice expressed DNA-PKcs while (D-F) were calculated from DNA-PKcs^PTKO^ kidney sections on NF or HF diet were treated with vehicle (saline) or cisplatin (15 mg/kg IP). (G-I) DNA-PKcs^PTKO^;Trip13^ΔStop^ kidneys were compared with or without cisplatin (**J**) Western blot analysis was performed for p-MLKL and total MLKL using protein lysates from the kidneys of various genotyped mouse groups. β-actin was used as a loading control. V-NF = vehicle-treated normal fat mice; C-NF = cisplatin-treated normal fat; V-HF = vehicle-treated high fat mice; C-HF = cisplatin-treated high fat mice; DNA-PKcs^PTKO^;Trip13^ΔStop^ over-expressing Trip13 = proximal tubule knockout of DNA-PKcs; ^ΔStop^ = Trip13 over-expression in proximal tubule; DNA-PKcs^PTKO^;Trip13^ΔStop^ = proximal tubule knockout of DNA-PKcs and over-expression of TRIP13.*P<0.05, **P<0.01, ****P<0.0001 significant difference between indicated groups.

Since Ki-67 is more broadly expressed during multiple phases of the cell cycle, including S, G2 and M [30], we investigated the number of phospho-Histone H3 (Ser10)-positive (denoted as pH3+) cells in the kidney sections obtained from the cisplatin-treated NF (0.4 ± 0.1%; n=8) and HF (0.2 ± 0.1%; n=5)-fed DNA-PKcs^CTRL^ mice (**Fig. 3B** and **3D**). No significant difference in the pH3 (Ser10) was detected in the DNA-PKcs^PTKO^ mouse kidneys under basal conditions during NF (0.30 ± 0.2%; n=3) or HF states (0.1 ± 0.1%; n=3) (**Fig. 3E**). Upon treatment with cisplatin, pH3+ nuclei were neither markedly changed in DNA-PKcs^CTRL^ mice fed NF or HF (**Fig. 3B**), nor NF DNA-PKcs^PTKO^ mice treated with cisplatin (**Fig. 3E**).

In vehicle-treated DNA-PKcs^CTRL^ or DNA-PKcs^PTKO^;Trip13^Stop/Stop^ mice kidneys, Ki67+ nuclei comprised 2.4 ± 0.5% (n=6) and 4.3 ± 1.3% (n=4), respectively, of the total tubular epithelial cells (**Fig. 3G**). Upon treatment with cisplatin, Ki-67 was detected in 1.9 ± 0.3% (n=9) of the total tubular epithelia from DNA-PKcs^CTRL^ kidneys. In the kidneys from cisplatin-treated DNA-PK^PTKO^;Trip13^ΔStop^ mice, the Ki-67 positive nuclei remained at similar levels (2.2 ± 0.4% n=7) with the other groups.

### Effects on programmed cell death due to DNA-PKcs

Apoptosis as detected by cleaved caspase-3 positive (CC3+) cells was either not (0.05 ± 0.02%; n=8) or rarely (0.03 ± 0.03%; n=6) detected in kidneys from NF or HF-fed DNA-PKcs^CTRL^ mice, respectively (**Fig. 3C** and **3F**). In the 7-8 hour water restricted mice treated with cisplatin, the CC3+ cells was increased in the kidneys from NF- and HF-fed DNA-PKcs^fl/fl^ mice to 0.33 ± 0.13% (n=7) and 0.15 ± 0.03% (n=6), respectively. In the vehicle-treated DNA-PK^PTKO^ mice fed NF- and HF had similarly low percentages of CC3+ cells at 0.08 ± 0.03% (n=4) and 0.03 ± 0.03% (n=3), respectively. Treatment with cisplatin in the NF-fed DNA-PK^PTKO^ mice significantly increased by ∼5-fold to 0.38 ± 0.16% CC3+ cells. In the more prolonged water restricted NF-fed DNA-PKcs^CTRL^ mice prior to injection with vehicle, the CC3+ cells remained low at 0.10 ± 0.05% (n=6), but the administration of cisplatin led to a nearly 30-fold increase to 2.9 ± 0.4% (n=9) (**Fig. 3I**). In NF-fed DNA-PK^PTKO^;Trip13^ΔStop^ mice, there was no detectable CC3+ cells in the vehicle-treated group. Following cisplatin treatment, the percentage was increased to 0.49 ± 0.17% (n=7), but was ∼6-fold lower than kidneys obtained from the mice that were water restricted for 7-8 hours (**Fig. 3I**).

To study an alternate mode of cell death in the kidney, a key executioner protein for necroptosis, MLKL, was examined by Western blot analysis (**Fig. 3J**). Our findings observed that there was an increased phosphorylation at Serine 345 in the cisplatin-compared to vehicle DNA-PKcs^CTRL^ mice kidneys. Moreover, the DNA-PK^PTKO^ mouse kidneys had augmented p-MLKL (Ser345) levels compared to comparable cisplatin-treated DNA-PKcs^CTRL^ kidneys (**Fig. 3J**).

### Role of DNA-PKcs on kidney injury in a mouse model of ischemia-induced AKI

In a second rodent model of AKI, we performed a surgical procedure in which both kidneys underwent short-term ischemia after which the kidneys were allowed to reperfuse for a period of 24 hours. In the PKcs^CTRL^mice there was a markedly higher level of serum creatinine (0.56 ± 0.18 mg/dL; n=5) compared to the sham-operated mice (0.10 ± 0.01 mg/dL; n=5) (**Fig. 4A**). In the DNA-PKcs^PTKO^ mice, the serum creatinine was significantly more elevated to 0.93 ± 0.37 mg/dL (P<0.05; n=4) compared to the sham group. There was a tendency for the tissue KIM-1 levels to increase in the IRI-versus sham-treated mouse kidneys, but it was not significant (**Fig. 4B**). Consistent with the changes in serum creatinine, the tubular injury in the DNA-PKcs^PTKO^ kidneys was significantly higher than the sham- (P<0.001) and IRI-treated (P<0.05) kidneys with the intact DNA-PKcs gene (**Fig. 4C-4F**).

**Figure 4.**
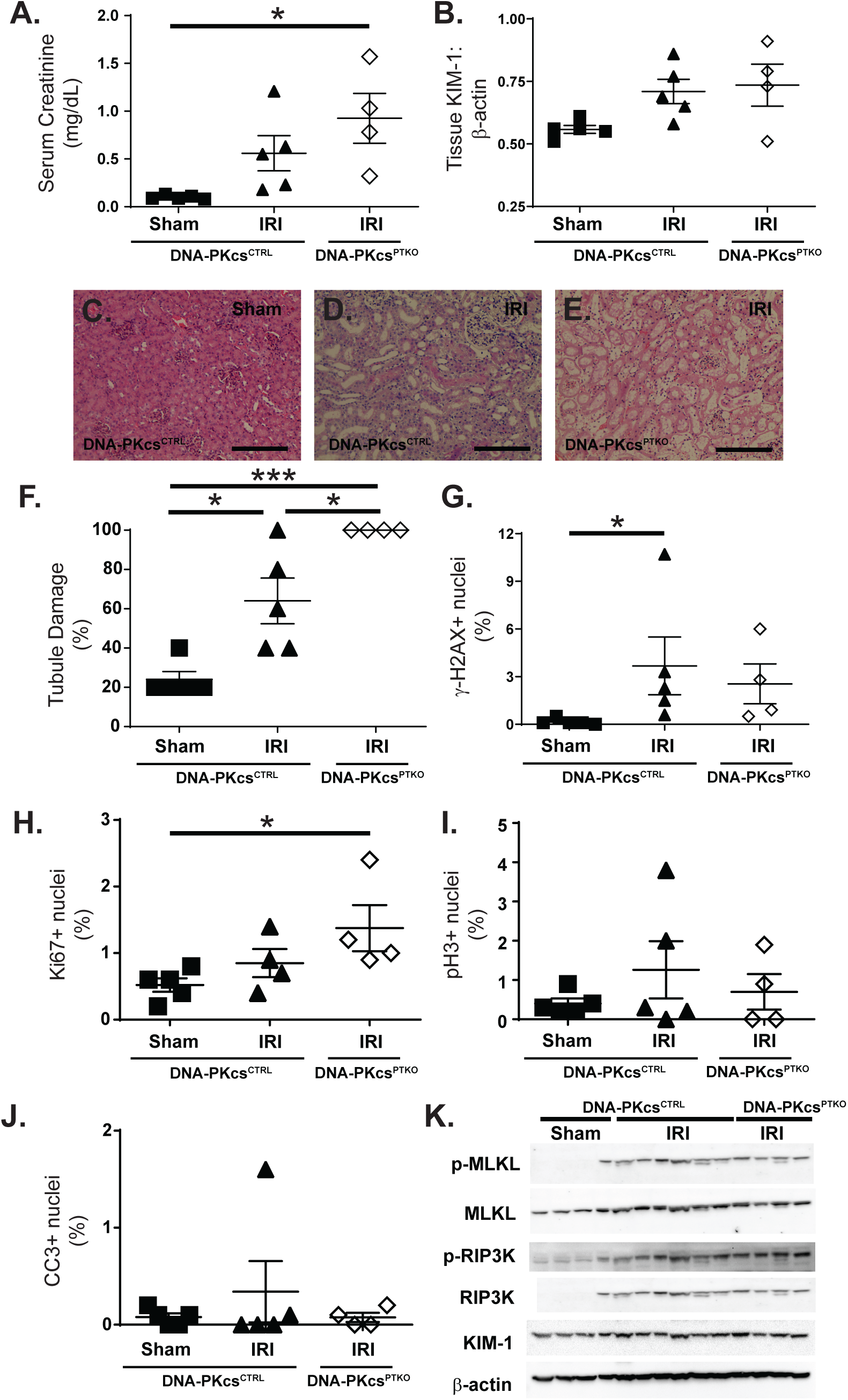
Selective loss of DNA-PKcs in the proximal tubules augments tubular damage following bilateral renal ischemia-reperfusion injury (bIRI). (A) Serum creatinine was measured after 24 hours following reperfusion. (B) Tissue changes in KIM-1 as determined by Western blot analysis. The blot for KIM-1 used for this analysis is shown in Fig. 4J. (C-E) Representative kidney sections from sham (C) or bilateral IRI (D, E) from DNA-PKcs^CTRL^ (C, D) or DNA-PKcs^PTKO^ (E) mice. (F) Tubular damage as a percentage of the total tubules counted was graphed from each individual mouse. *P<0.05, ***P<0.001 significant difference between indicated groups. (G) DNA damage (γ-H2AX), (H) proliferation (Ki-67), (I) mitotic activity (p-H3 (Ser10)), (J) apoptotic cell death (cleaved caspase-3) was determined by counting marker-specific nuclei relative to the total population of nuclei (hematoxylin). (K) Western blot analysis of kidney lysates for nephrotic pathways (p-RIP3K and p-MLKL) and KIM-1 (tissue injury marker shown in Fig. 4B) in sham and IRI-injured kidneys. β-actin was used as a loading control. IRI = bilateral ischemia-reperfusion injury

Unlike the cisplatin treatment group, there was no obvious differences in phenotypic markers that we assessed in this set of kidney samples. DNA damage was nearly absent in sham-operated kidneys (0.2 ± 0.1%; n=5), and showed a mild, but significantly elevation in the γH2AX-positive nuclei in the IRI-treated DNA-PKcs^CTRL^ mouse kidneys (3.7 ± 1.6%; n=5) (**Fig. 4G**). In the DNA-PKcs^PTKO^ kidneys, there was also a markedly higher number of γ-H2AX positive nuclei (2.6 ± 1.3%; n=4) compared to the sham-treated mouse kidneys, but not different from the IRI-treated DNA-PKcs^CTRL^ mice. The proliferation status of sham-operated kidneys was minimal as determined by the Ki-67 positive nuclei in **Fig. 4H** (0.5 ± 0.%; n=5). The bilateral IRI kidneys were mildly increased to (0.9 ± 0.2%; n=4), but did significantly increase by ∼3-fold to 1.4 ± 0.7% (n=4) in the DNA-PKcs^PTKO^ kidneys. There were no significant changes in the number of cells entering mitosis (pH3+; **Fig. 4I**) or apoptotic cell death (CC3+; **Fig. 4J**) regardless of the DNA-PKcs status in the proximal tubules following IRI.

Instead, necroptotic cell death pathways (p-MLKL and RIP3K) were markedly increased after bilateral IRI (**Fig. 4K**). Similar increases in p-MLKL and RIP3K was observed in the DNA-PKcs^PTKO^ kidney lysates compared to sham-operated kidneys.

### Role of DNA-PKcs on kidney fibrosis and kidney injury in a mouse model of unilateral ureteral obstruction

UUO is a classic model of progressive kidney fibrosis and chronic kidney disease (CKD). Male and female mice exhibited similar effects on biological changes to the kidney architecture and markers for DNA damage and cell cycle, so the data from both sexes using this model were combined for phenotyping (**Fig. 5** and **6**).

**Figure 5.** Effect of a proximal tubule-specific loss in DNA-PKcs on biological parameters in the kidney following unilateral ureteral obstruction. (A-D) Representative images from sham (A) or UUO kidneys with (B) DNA-PKcs^CTRL^ (C) DNA^PTKO^, and (D) DNA^PTKO^;Trip13^ΔStop^ genotypes. DNA^PTKO^ is the same as DNA-PKcs^PTKO^ strain, but shortened in this figure to fit into the graph. (E) Serum creatinine was measured at the end of the 7 day experimental period. Kidney sections were stained and quantitatively analyzed by counting antibody-specific staining nuclei to determine: (F) DNA damage (γ-H2AX+); (G) cell proliferation (Ki-67+); (H) Mitotic activity (pH3+), and (I) apoptosis (CC3). * P<0.05, ** P<0.01, ***P<0.001, **** P<0.0001 indicates significant differences between groups. pH3 = phospho-Histone H3 (Ser10); CC3 = cleaved caspase-3. Scale bar = 200 µm.

**Figure 6.**
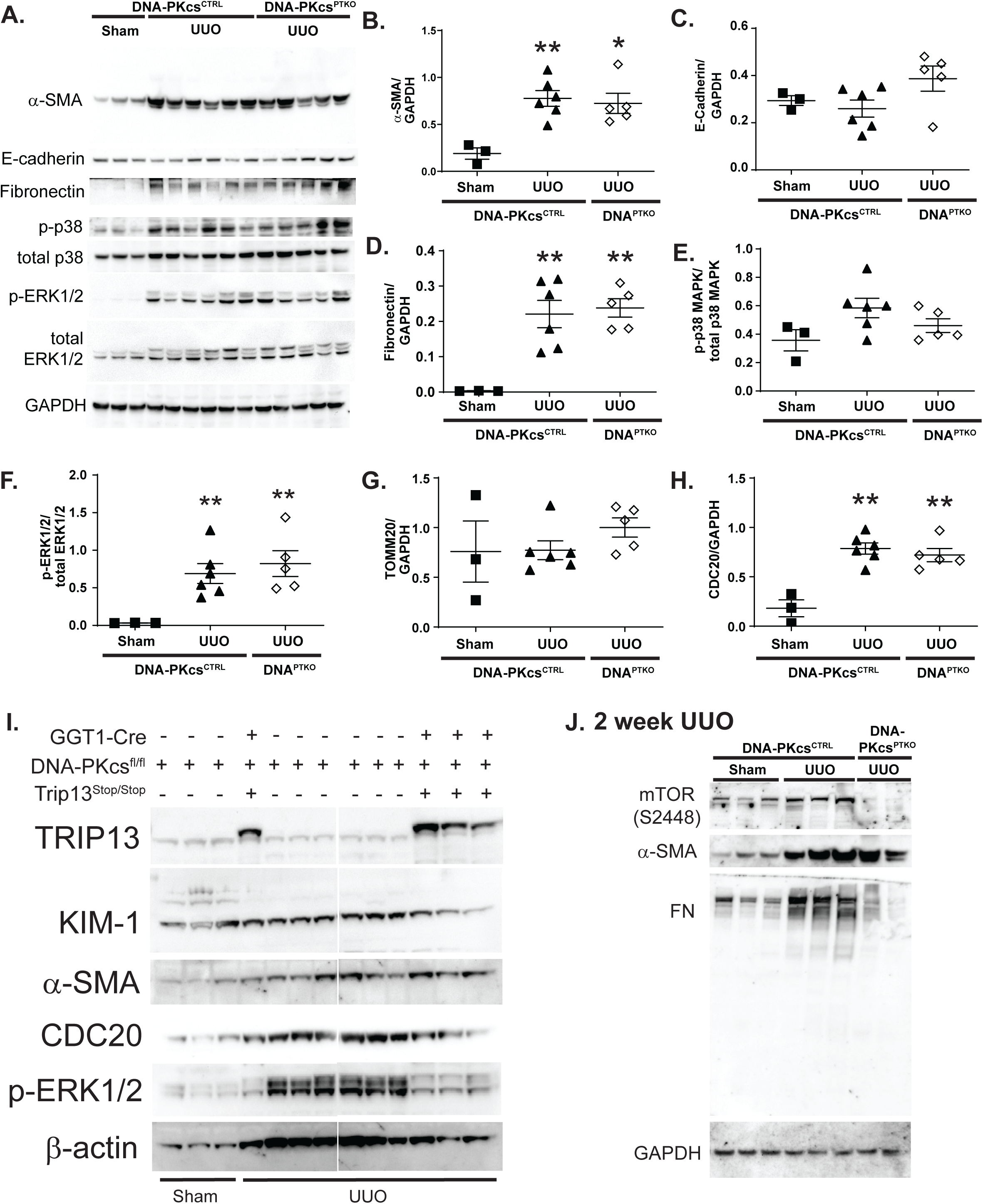
Effect of the absence of DNA-PKcs in the proximal tubules on fibrotic marker production and MAPK signaling following UUO. (A) Representative Western blot images using protein lysates from the sham and UUO kidneys are shown for day 7 (A, I) and 14 (J) after the start of the UUO procedure. In this figure, primary antibodies were used to target: α-SMA, fibronectin, E-cadherin, p-p38 MAPK, total p38 MAPK, p-ERK1/2, and total ERK1/2. GAPDH was used as a loading control. DNA^PTKO^ is the same as DNA-PKcs^PTKO^ strain, but shortened in this figure to fit into the figure. (B-H) Densitometry of the bands was quantitated using ImageJ for: (B) α-SMA, (C) E-Cadherin, (D) fibronectin, (E) phospho-to-total p38 MAPK, (F) phospho-to-total ERK1/2, (G) CDC20. * P<0.05, ** P<0.01 indicates significant differences between sham group. (H) Sirius red staining of kidney sections from the sham and UUO groups. (I) Additional Western blots were performed to examine protein levels for: KIM-1, CDC20, α-SMA, TRIP13 and p-ERK1/2 using kidney lysates from sham and UUO-treated kidneys after 7 days. β-actin was used as a loading control. A break was made in the membrane due to an aberrant sample in the gel. (J) Representative blots were shown for mTOR (S2448), α-SMA, FN and GAPDH using kidney lysates from DNA-PKcs^CTRL^ and DNA-PKcs^PTKO^ mice after 2 weeks from the start of the sham or UUO procedure.

As expected, sham-treated exhibit no histological injury (**Fig. 5A**). However, there were substantial architectural changes in the UUO kidneys, such as the dilation of renal tubules. For male and female mice with normal endogenous expression of DNA-PKcs, ureteral ligation did not exhibit any visual changes in the kidney histology between the UUO-treated groups (**Figure 5B-5D**). There was no significant changes in the circulating levels of creatinine (0.10 ± 0.01 mg/dL; n=13) between UUO compared to sham-operated DNA-PKcs^CTRL^ mice (0.08 ± 0.01 mg/dL; n=8) due to the ability of the contralateral kidneys to continue excreting creatinine. In the DNA-PKcs^PTKO^ mice, the serum creatinine was similarly non-significantly higher (0.15 ± 0.04 mg/dL; n=8) compared to the other mouse groups (**Fig. 5E**).

The extent of DNA damage as determined by γ-H2AX positive nuclei in UUO DNA-PKcs^CTRL^ kidneys (2.4 ± 0.4%; n=13) was significantly higher (P<0.05) compared to kidneys from sham-operated DNA-PKcs^CTRL^ mice (0.3 ± 0.1%; n=7) (**Fig. 5F**). DNA-PKcs^PTKO^ kidneys did not affect the magnitude in γ-H2AX positive nuclei (2.5 ± 0.5%; n=8) compared to UUO DNA-PKcs^CTRL^ mouse kidneys. The sham-operated DNA-PKcs^CTRL^ mice at both days 7 (1.1 ± 0.4%; n=6) and 14 (1.0 ± 0.1; n=6) after surgery exhibited minimal proliferation as detected by the low percentage of Ki67-positive nuclei (**Fig. 5G**). Similarly, the mitotic marker, p-Histone H3 (Ser10) or pH3+, was rarely detected in sham-operated DNA-PKcs^CTRL^ mouse kidney sections (0.09 ± 0.03%; n=8) (**Fig. 5H**). Obstruction of the ureter led to a time-dependent increase in Ki67-positive nuclei to 1.8 ± 0.2% at day 7 (n=13) (**Fig. 5G**) and 7.5 ± 1.6% (n=7) at day 14 (not shown). In DNA-PKcs^PTKO^ kidneys, Ki67-positive nuclei was observed at 2.5 ± 0.4% (n=8) at day 7 (**Fig. 5G**) in the UUO kidneys, which continued to increase through day 14 (5.6 ± 0.5%; n=3) (not shown). The cellular proliferation was not significantly different than DNA-PKcs^CTRL^ mouse UUO kidneys. A significantly higher percentage of pH3+ cells were detected in both UUO DNA-PKcs^CTRL^ (1.5 ± 0.2%; n=13) or DNA-PKcs^PTKO^ (1.4 ± 0.2%; n=8) kidneys (**Fig. 5H**). Apoptotic cell death was not detected in the sham-treated kidneys, but there was detectable increase in CC3+ cells from UUO DNA-PKcs^CTRL^ and DNA-PKcs^PTKO^ kidneys to 1.1 ± 0.1% (n=11) and 0.8 ± 0.2% (n=6), respectively (**Fig. 5I**).

Using kidney protein lysates, we detected significantly elevated KIM-1 levels (P<0.05), which is a marker of tubular damage, by Western blot in the UUO compared to sham DNA-PKcs^CTRL^ kidneys (**Fig. 6I**), but there was no significant difference in KIM-1 levels between the DNA-PKcs^PTKO^ versus DNA-PKcs^CTRL^ UUO kidneys. Fibrotic markers, α-smooth muscle actin (α-SMA) (**Fig. 6A** and **6B**), and fibronectin (FN) (**Fig. 6A** and **6D**), were significantly elevated in UUO compared to sham DNA-PKcs^CTRL^ kidneys, but no additional change was detected in the DNA-PKcs^PTKO^ kidneys. E-cadherin was not changed in any of the conditions or genotypes tested (**Fig. 6A** and **6C**).

During UUO, activated p38 MAPK (**Fig. 6E**) and ERK1/2 (**Fig. 6F**) signaling were increased compared to sham conditions (**Fig. 6A**). At day 7, the UUO-treated DNA-PKcs^PTKO^ kidneys did not exhibit any difference in either MAPK activity compared to the DNA-PKcs^CTRL^ kidneys. CDC20 is a M phase marker, which was significantly increased (**Fig. 6G**) in UUO compared to sham DNA-PKcs^CTRL^ kidneys, but was not affected in the absence of DNA-PKcs following UUO. Sirius red staining was observed to be elevated in UUO DNA-PKcs^CTRL^ kidneys with no significant difference in the DNA-PKcs^PTKO^ kidneys (**Fig. 6H**).

### TRIP13 over-expression reduces tubular remodeling and interstitial fibrosis

Prior work by Banerjee et al. [15] in head and neck cancer cells demonstrated that TRIP13 promoted DNA repair through NHEJ by direct binding to DNA-PKcs. Subsequently, numerous groups have demonstrated that abnormal levels of TRIP13 expression are associated with the progression of different types of cancer [31–38]. In our lab, we investigated the effect of a proximal tubule-specific TRIP13 over-expression (ΔStop) [4] by breeding into the DNA-PKcs^PTKO^ mouse to determine the effect on DNA damage during UUO and its subsequent role, if any, in tubular remodeling and fibrosis progression. No significant difference was detected in the DNA damage index (1.7 ± 0.8%; n=4) in the DNA-PKcs^PTKO^;Trip13^ΔStop^ kidneys compared to the UUO-treated DNA-PKcs^PTKO^ kidneys as mentioned earlier in **Fig. 5F**. Of interest, however, was a significantly higher proliferation status (11.6 ± 0.5% Ki67+; n=4) and mitotic index (2.5 ± 0.2% pH3+; n=4) in the TRIP13-overexpressing (DNA-PKcs^PTKO^; Trip13^ΔStop^) compared to the DNA-PKcs^PTKO^ kidneys (**Fig. 5H**). Apoptotic cell death was not significantly different in the TRIP13-expressing (DNA-PKcs^PTKO^;Trip13^ΔStop^) (0.85 ± 0.12%; n=3) UUO kidneys compared to the values from the DNA-PKcs^PTKO^ (0.82 ± 0.21%; n=7) or DNA-PKcs^CTRL^ (0.96 ± 0.6%; n=9) UUO kidneys. Apoptotic cell death was slightly lower, but non-significantly to 0.6 ± 0.1% CC3+ cells in the UUO DNA-PKcs^PTKO^; Trip13^ΔStop^ kidneys. By Western blot analysis, the TRIP13-overexpression in the UUO DNA-PKcs^PTKO^; Trip13^ΔStop^ kidneys showed a reduced production of KIM-1 (**Fig. 6J**) with a concomitantly lower activation of ERK1/2.

### Single Nuclei RNAseq demonstrated attenuation of proximal tubule injury in DNA-PKcs^PTKO^; Trip13^ΔStop^

In total, there were 62,059 (59,769 after quality control) cells isolated from kidney and sequenced across the 4 groups, including vehicle-treated DNA-PKcs^CTRL^ (V-NF), cisplatin-treated DNA-PKcs^CTRL^ (C-NF), vehicle-treated DNA-PKcs^PTKO^; Trip13^ΔStop^ (V-NF) and cisplatin-treated DNA-PKcs^PTKO^; Trip13^ΔStop^ (C-NF) (**Fig. 7**). Clustering analysis using known canonical markers for kidney cell types identified 16 distinct cell clusters consisting of a small number of cells to as many as 8,748 cells per cluster in the proximal tubules group (**Fig. 7A** and **7B**). The distribution, number of cells, and clustering of identified cell types were consistent across all 4 groups (**Fig. 7C, Table SX-ClusterCount**). Along with the expected cell types, a unique cluster expressing high levels of Havcr1 (Kim-1), a tubule specific injury marker suggesting a subset of proximal tubules with severe injury. Since cisplatin is a known to accumulate and promote proximal tubule toxicity and injury, differential expression analysis was focused on proximal tubule cells (**Table SX-DEGPTXGroup**).

**Figure 7.**
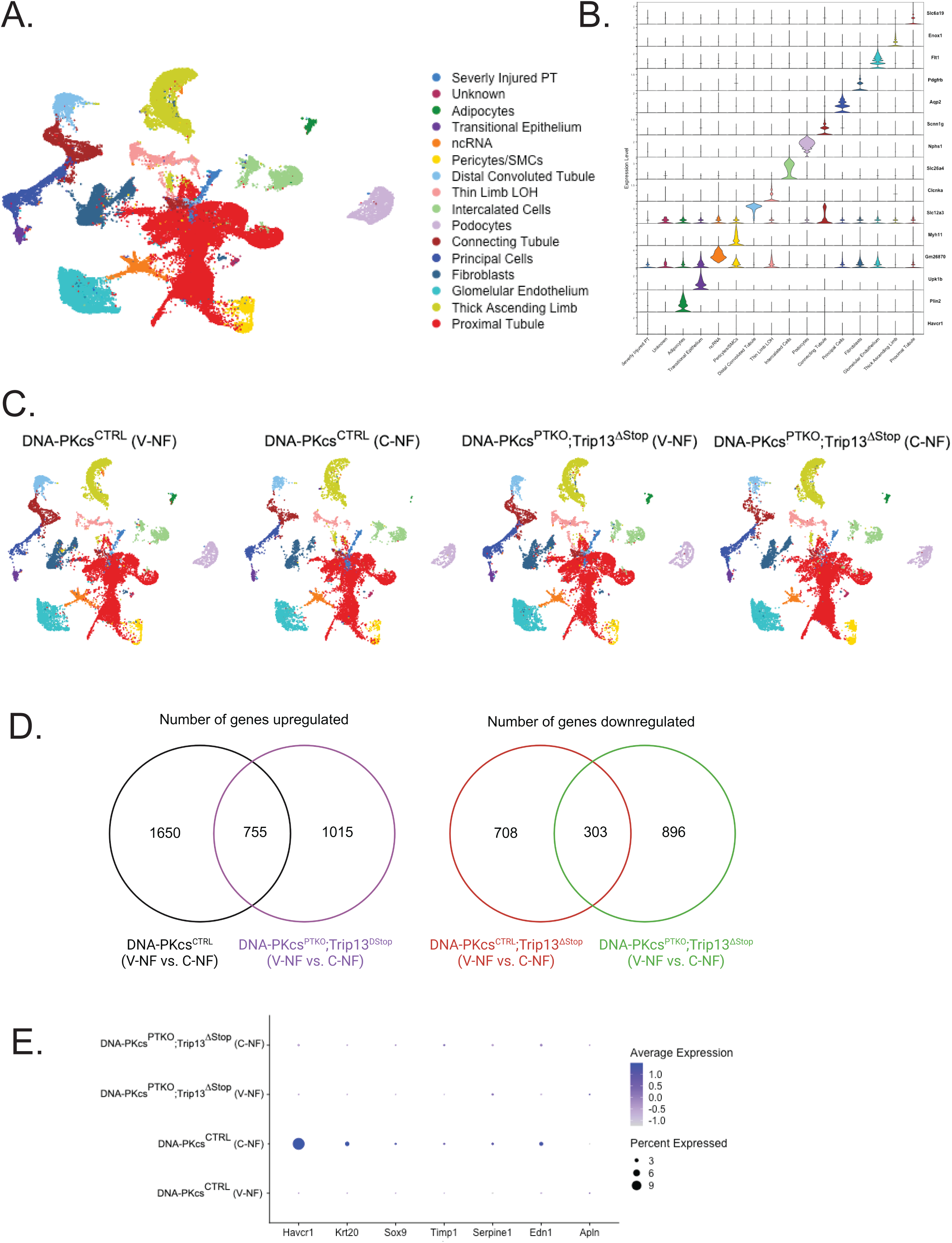
Single nuclei RNASeq analysis and informatics. (A) UMAP of the single nuclei RNASeq analysis is shown from 4 different kidney treatment groups. Cells are clustered in 2 dimensions using the UMAP dimensionality reduction technique, followed by integration with harmony and annotated by cluster labels. 16 different cell types were identified and annotated, as shown on the right side. (B) Violin plots depicting the gene expression level (depicted on the vertical left axis) for each of the canonical markers (depicted on the right vertical axis) used for cell type annotation across the 16 clusters. (C) UMAP plots split by the 4 groups. (D) Venn diagrams of overlap of up and downregulated genes for each group comparison (V vs C). (E) Dot plot displaying injury related markers and their up/down regulation in the 4 groups.

There were a large number of differentially expressed genes (DEG) between vehicle-treated DNA-PKcs^CTRL^ (V-NF) vs cisplatin-treated DNA-PKcs^CTRL^ (C-NF), including 2,405 upregulated and 1,011 downregulated (total DEG = 3,416 at Log2FC>±0.6 and adjusted p<0.05), suggesting that the single dose treatment of cisplatin resulted in severe injury and a significant disruption to the whole transcriptome. In comparing cisplatin-treated DNA-PKcs^PTKO^; Trip13^ΔStop^ (C-NF) with its control [vehicle-treated DNA-PKcs^PTKO^; Trip13^ΔStop^) there were 15% less DEG (1,770 upregulated and 1,199 downregulated (total DEG = 2969 at Log2FC>±0.6 and adjusted p<0.05) compared to DNA-PKcs^CTRL^ (V-NF vs C-NF) comparison suggesting that the cisplatin injury was somewhat attenuated. Comparing upregulated and downregulated DEG between each comparison, there was an overlap of core DEG (upregulated=755 and downregulated= 303, **Table SX**) associated with cisplatin treatment regardless of the strain i.e., DNA-PKcs^CTRL^ or DNA-PKcs^PTKO^; Trip13^ΔStop^ (**Fig. 7E**). While there was a core group of DEG similarly dysregulated, most DEG were unique to either cisplatin treated DNA-PKcs^CTRL^ (compared to vehicle) or cisplatin-treated DNA-PKcs^PTKO^; Trip13^ΔStop^ (compared to vehicle control). Top pathways overrepresented in core DEG using Reactome pathway analysis identified no enrichment for upregulated genes (n=755). Moreover, no statistically significant (FDR<0.05) pathway/enrichment in DEG was observed in either group comparison for upregulated genes (n=1,650 or 1,015, respectively), although using unadjusted p-value (<0.05) several pathways (nuclear receptor transcriptome, Rho GTPase, and RNA polymerase III chain elongation) were identified for unique genes observed in DNA-PKcs^PTKO^; Trip13^ΔStop^ (V-NF vs C-NF).

Top pathways overrepresented in core downregulated DEG (n=303) identified enrichment for pathways associated with peptide chain elongation, translation elongation, nonsense mediated decay. For unique set of down-regulated DEG in the DNA-PKcs^PTKO^; Trip13^ΔStop^ (V-NF vs C-NF), top pathways identified were associated with respiratory electron transport, complex 1 biogenesis, mitochondrial protein degradation). An evaluation of Havcr1 (Kim-1) and other associated proximal tubule injury markers showed that the highest number of injured cells with correspondingly highest expression (**Fig. 7E**) was measured in the DNA-PKcs^CTRL^ (C-NF) mouse strain. In contrast, cisplatin treated DNA-PKcs^PTKO^; Trip13^ΔStop^ (C-NF) had comparably lower number of injured cells and gene expression levels compared to either of the vehicle treated groups. The genetic profiles of the proximal tubules demonstrating reduced changes in gene expression levels due to a lesser cellular damage would be consistent with the phenotypic observations in the mice kidneys expression proximal tubular TRIP13 even with the genetic removal of DNA-PKcs.

### Inhibition of DNA-PKcs in proximal tubular epithelial cells decreases cell number *in vitro*

Incubation of mouse TKPTS proximal tubule cells with a highly selective DNA-PKcs inhibitor, NU7441, demonstrated a modest dose-dependent (0, 0.1 and 0.2 µM) reduction in cell numbers (**Fig. 8A**). At the higher dose, the cell numbers were ∼15% lower (239,739 +/- 11,444 cells; n=4) compared to the time-control (DMF) group (297,400 +/- 9,672 cells; n=3). In a separate series of experiments, TKPTS cells were synchronized using a double thymidine block protocol in the presence of either a highly selective DNA-PKcs inhibitor, NU7441, or vehicle (DMF) during the second thymidine block period. Afterwards, the cells were maintained in normal DMEM:F12 media with 5% FBS for up to 16 hours. Propidium iodide staining was performed on the isolated cells and sorted by flow cytometry (**Fig. 8B**). NU-7441 slowed the progression through the cell cycle compared to the control (vehicle) treated cells. In 2 hours, the TKPTS cells have progressed from primarily G1/S to G2 whereas the NU7441 was not proceeding to G2 until about 5 hours after release from NU7441. By 16 hours, the NU7441 cells have a similar profile as the vehicle-treated cells, but there was still a higher percentage of cells at the G2/M phase. The reduced number of cells in the presence of NU7441 did not appear to be associated with cell death, but rather a slower progression through the cell cycle.

**Figure 8.**
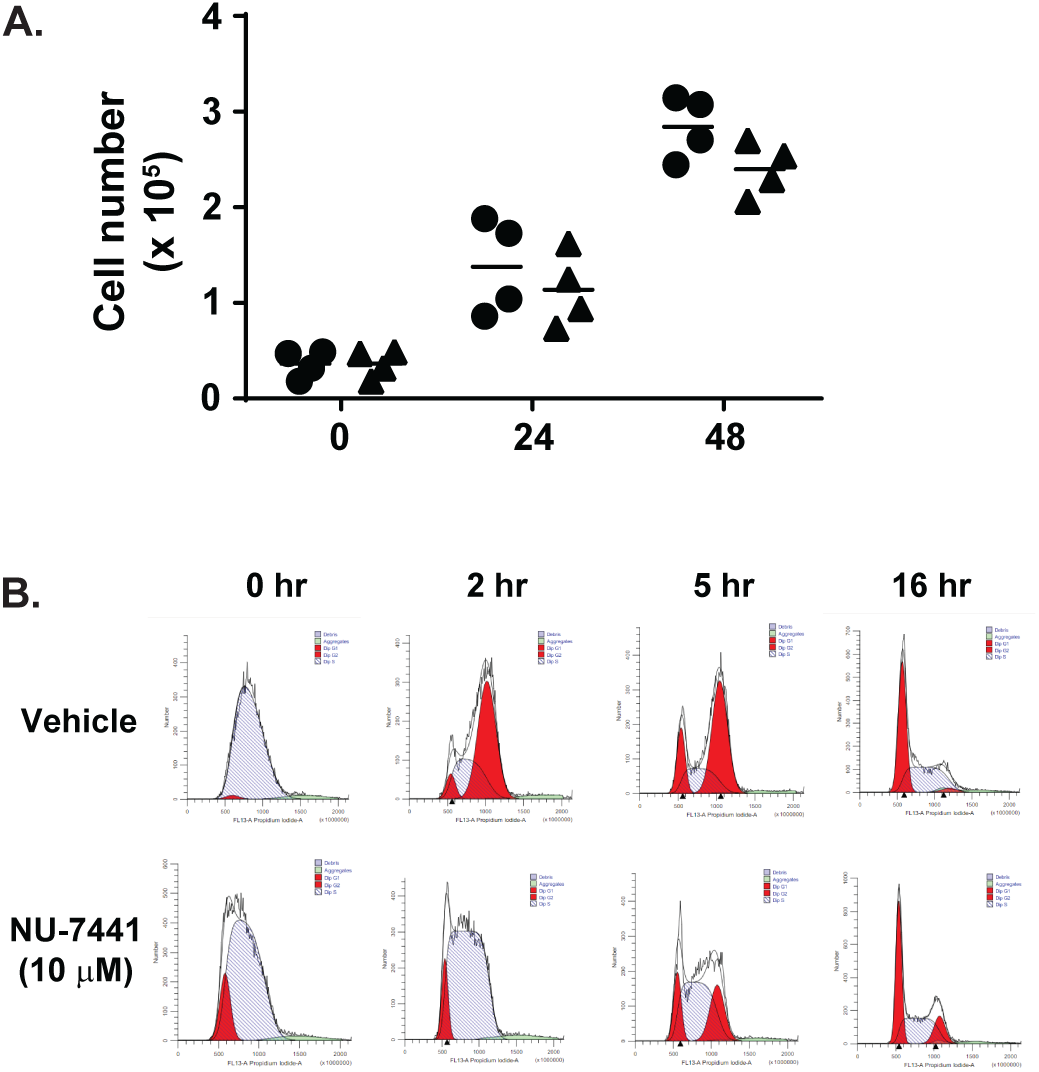
Effect of DNA-PKcs inhibition on cell function in vitro. (A) Cell numbers of TKPTS cells were counted by hemocytometry over a 48 hour period in the presence or absence of NU-7441 (10 μM). Vehicle = dimethylformamaide (DMF). λ= vehicle-treated cells; π = NU-7441 treated cells. 3 different experiments were performed and averaged for this graph. (B) Cell cycle analysis was performed in TKPTS cells synchronized by double thymidine block in the presence or absence of NU-7441 during the second thymidine block period. At time 0 and subsequent time points, cells were harvested and stained with propidium iodide for cell cycle analysis. A representative experiment is shown in this figure. 4 separate experiments were performed with similar profiles.

## Discussion

The catalytic subunit of DNA protein kinase (DNA-PKcs) plays a crucial role in a multitude of biological pathways in the mammalian system from sensing DNA damage, activating DNA repair pathways, and regulating cellular metabolism [39]. In the present study, we demonstrated that the loss of DNA-PKcs in the proximal tubules (DNA-PKcs^PTKO^) can exacerbate tubular DNA damage and cell injury due to exposure of two different types of acute kidney injury (AKI). This is in confirmation of other recent studies whereby DNA damage albeit variable is an early event following AKI and plays a fundamentally important role in the pathogenic changes in the renal tubules [4, 5, 10]. DNA-PKcs is a member of the PI3KK family along with ATR and ATM [40], in which they present atypical kinase activity during various types of genome damage. In certain situations, DNA-PKcs can work together with either ATR and ATM to phosphorylate proteins involved in the DNA damage checkpoint. Unlike ATR, which has been established to be activated by ssDNA [41], DNA-PKcs is activated by the exposure of the ends from double stranded DNA breaks (DSBs) and functions primarily through the non-homologous end joining (NHEJ) pathway [10]. Similar to proximal tubule-specific ATR knockout mice, there was no obvious functional or histological differences in the DNA-PKcs^PTKO^ mouse compared to their genetic control kidneys (DNA-PKcs^CTRL^). The absence of ATR in the proximal tubules demonstrated increased DNA damage, apoptosis and tubular damage in two different forms of AKI [5], which complements the results in our study. The volume status of the mice or the fat diet made the mice more susceptible to the cisplatin effects in the absence of the DNA-PKcs in the proximal tubules. The increased susceptibility towards kidney damage by cisplatin exposure due to a short-term switch to a high fat diet was similarly observed in a recent study by Kim et al. [37]. Proliferation was not markedly different in the kidneys after cisplatin treatment similar to findings by Kishi et al. [5] in their PT ATR KO mouse, but the cell death was higher in the DNA-PKcs knockout kidneys.

The increased DNA damage is likely to have preceded the tubular injury associated with pharmacological inhibition using NU-7441 as recently shown by our lab [4]. NU-7441 appeared to have a more rapid and devastating effect on the mouse kidney due to cisplatin exposure compared to the genetic knockout mouse model. One obvious possibility is that the biological role for DNA-PKcs is essential for cell function outside of the kidney, so the systemic inhibition by NU-7441 may only be a part of the overall cause for the morbidity and the unexpected mortality of some mice. Another reason is the differences observed on the function of DNA-PKcs using loss of function versus genetic deletion systems. In a recent study by Menolfi et al. [36], expression of a loss-of-function ATR variant exhibited prolonged binding to the ssDNA end that would block downstream DNA repair systems [36]. Further review of the PI3KK family suggested that similar differences in the type of functional loss of the protein may determine distinct outcomes of the DNA damage [40]. The binding of pharmacologically inhibited DNA-PKcs onto the exposed DNA ends may prevent the entry of protein complexes needed for the completion of the end ligation step [40]. In our present study, the absence of the DNA-PKcs expression in the proximal tubules would not prevent the end ligation block at that step and ligation may be able to occur, albeit less efficiently to repair the breaks through ATM and homologous recombination. In either situation, however, both the inhibition of DNA-PKcs using NU-7441 or the genetic approach to delete DNA-PKcs from the proximal tubules led to an increased level of tubular injury and DNA damage following cisplatin treatment.

In our second model of AKI, bilateral renal IRI, the DNA-PKcs^PTKO^ mice showed increased serum levels of creatinine following IRI compared to DNA-PKcs^CTRL^ mice, but the DNA damage and cell proliferation (Ki67 and p-pH3) were not markedly different between genetic models with or without proximal tubular expression of DNA-PKcs. Of note, the overall the percentage of tubular cells with DNA damage was considerably lower than observed with cisplatin, which is consistent with prior observations in other labs [5]. In this particular model of AKI, it is possible that the role of DNA-PKcs has other pleiotropic functions outside of maintaining genome integrity to control nutrient and energy availability as previously shown in the skeletal muscle [17]. More specifically, DNA-PKcs acts to negatively reduce mitochondrial function through an inability to efficiently activate the HSP90α/AMPK pathway. This can lead to increased susceptibility of the cells towards metabolic changes that will reduce their capacity to protect themselves [17]. In this scenario, one would anticipate that reduced DNA-PKcs would facilitate a protective effect in the kidney during ischemia-reperfusion where the cells would be suffering from a lack of energy production needed for regeneration of the damaged cells. Alternatively, there is prior work demonstrating that DNA-PKcs can protect HIF-1α from degradation to promote cell survival, which can occur in the absence of double stranded DNA breaks [42]. Other PI3KK family members, including ATM, which senses DSB [43], and ATR, which is activated in response to single stranded DNA breaks and replication stress [44], can work cooperatively to regulate HIF-1 biosynthesis upon hypoxia. The role of DNA-PKcs and how its absence can affect cellular survival is likely context dependent in terms of the ability of the cell to adapt to changes in its internal milieu due to exposure of various types of damaging stimuli.

In a recent study where ATR was deleted from the proximal tubule, there was no difference in the extent of the kidney dysfunction early after the ischemia-reperfusion step, but instead there was a markedly slower recovery of the kidney with considerable inducement of fibrosis over a period of 3 weeks [40]. During UUO, the percentage of γ-H2AX, which is used as a proxy for DNA damage, was considerably elevated in the ATR deleted versus ATR floxed mice [5]. In our study, however, the absence of DNA-PKcs during UUO did not affect the overall number of γ-H2AX positive nuclei. In addition, there was no significant difference in the levels of fibronectin or a-smooth muscle actin in the DNA-PKcs^PTKO^ mouse kidneys compared to the control UUO kidneys. These findings may suggest that the type of genomic DNA damage during UUO progression occurs through either replication stress or formation of single stranded DNA ends with a lesser emphasis on double stranded DNA breaks [5]. Alternatively, the repair of damaged DNA normally fixed by DNA-PKcs could be compensated by other pathways, albeit less efficiently, such as Mre11-Rad50-NBS1 complex [35].

Consistent with prior published work [5], there was increased cellular proliferation and mitotic progression in the UUO versus sham or contralateral kidneys. In the absence of DNA-PKcs in the proximal tubule during UUO, there was no difference compared to the normal DNA-PKcs-expressing kidneys. As an adaptor protein, TRIP13 has been shown to interact directly with DNA-PKcs in cancer cells to facilitate DNA repair through the non-homologous end joining (NHEJ) pathway [15]. Moreover, there are numerous studies that have demonstrated that abnormal levels of TRIP13 expression are associated with the progression of different types of cancer [31–38]. Although there was no obvious proliferative advantage in the TRIP13 over-expressing kidneys, TRIP13 appeared to help in protecting the kidneys from excessive DNA damage and cellular injury during cisplatin administration in this study and in our recent work [4]. In the absence of DNA-PKcs, over-expression of TRIP13 did reduce the DNA damaged exerted by cisplatin and progression towards apoptotic cell death. There is evidence that the necrotic pathway activation by cisplatin could be mitigated by TRIP13 over-expression even in the absence of DNA-PKcs. These findings would suggest that TRIP13 can function independently of DNA-PKcs during tubular epithelial cell injury to promote a protective effect on DNA damage. However, this may be context-dependent as the positive effects exerted by TRIP13 was not observed in UUO kidneys where there is persistent tubular injury due to the blockade of the ureter. Similar to other studies in the cancer field demonstrating that TRIP13 can promote active proliferation, the UUO kidney cells in our experiments showed a similarly elevated percentage of Ki67+ and pH3+ cells indicating a more active cell cycle status. However, there was no obvious difference in the size or number of dilated tubules in the kidneys regardless of TRIP13 over-expression, so further studies are needed to better understand the potentially distinct role of TRIP13 in both acute and chronic injury states of the kidney and how DNA-PKcs plays a role, if any, in the recovery of the damaged tubular epithelia.

Our results, however, differ from studies by Wang et al. [45] in which their lab generated a proximal tubular deletion of DNA-PKcs using the androgen-specific Kap2 promoter to drive Cre recombinase. One possible reason is that there may be strain differences between our mice from the NIH, which was on a C57Bl/6J background. The DNA-PKcs^fl/fl^ mouse strain used in our study was backcrossed to the C57Bl/6J mouse at least 10 generations by the Chung lab prior to our breeding protocols. However, there is a lack of information about the method and the genetic background of the DNA-PKcs^fl/fl^ in the original [46] and a more current publication by the Wang group [45]. Second, there were differences in the experimental protocols used between the studies: a) a higher dose of cisplatin (20 mg/kg IP) was used by Wang et al. [45] versus our present study (15 mg/kg IP); b) our study performed a water restriction protocol prior to the injection of cisplatin; and c) there was a considerably longer ischemia time (30 min) compared to our study (21.5 min). Their length of ischemia likely results in considerable necrosis that may lead to a differential activity through DNA-PKcs that is not observed in our protocol where we attempted to achieve sublethal injury. A third reason may be related to the interpretation of the results based on the Kap2 promoter activity in the kidney [45]. In their study, the authors stated that the results for male and female mice were combined as both sexes showed the same biological effect. This is somewhat contrary to what one would anticipate based on the initial study by Li et al. [47], who demonstrated that female mice exhibited extremely low to minimal Kap2 activity. Unlike males, the female mice with the Kap2 promoter would be expected to behave similar to non-Cre-expressing mice, so a mixture of the sexes into their analyses should have a broader range of biological outcomes. This was not the case where their findings were very narrow in their injury and subsequent response. These contradictory studies would suggest that further studies are needed to better clarify the role of DNA-PKcs in the normal and injured kidneys.

In summary, DNA-PKcs plays a fundamentally key role to mitigate tubular epithelial cell damage following exposure to damaging acute stimuli in the kidney. In the presence of constant stimuli that leads to a long-term change in tubular architecture, a loss in DNA-PKcs in the proximal tubule may not have a key role in cell recovery and affect the fibrotic progression in the kidney. This study demonstrates that DNA repair through the actions of DNA-PKcs is crucial to the normal and injury situations and needs to be considered during therapies that are focusing on the inhibition of this protein as it might impact the normal functioning of the kidney.

## Supporting information

Cluster Counts

DEG

## Acknowledgements

F.P. conceptualized and designed the experiments., M.S., J.Y., D.M.C., K.R.R., J. C., M.R.G, L.C., and F.P. provided reagents and performed the experiments, L.C., M.R.G., F.P., M.S. analyzed the data, M.S., L.C., M.R.G., and F.P. wrote and edited the manuscript.

The work performed through the UMMC Molecular and Genomics Facility is supported, in part, by funds from the NIGMS, including the Molecular Center of Health and Disease (P20GM144041-M.R.G), Mississippi INBRE (P20GM103476) and Obesity, Cardiorenal and Metabolic Diseases-COBRE (P30GM149404).

---

**Supplemental Figure 1.**
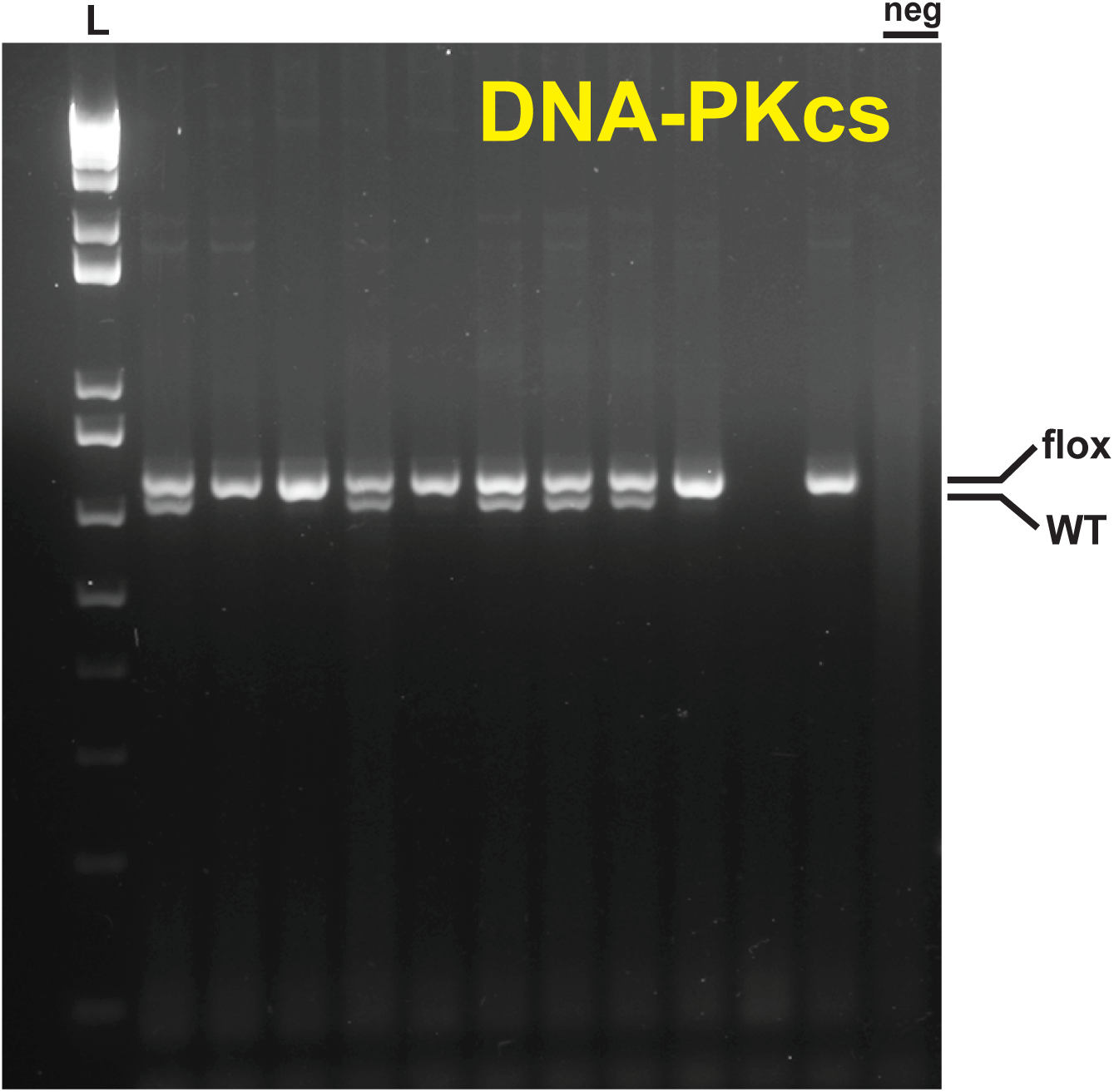
PCR genotyping of DNA-PKcs. As described in the Methods section, specific PCRprimers were used as described by Mishra et al. (Nature Communications, 2015) from the Chung lab. Standard PCRreagents and conditions were used and run on a 2% agarose gel with ethidium bromide to resolve the distinct PCRproduct bands between the wild-type and floxed DNA sequences. Each of the lanes are individual genomic DNA samples from multiple litters of mice. flox = 748 bp PCR product; WT= wild-type 706 bp PCRproduct. L = 1 kb+ DNA ladder (Invitrogen), neg = negative control (no genomic DNA template, but has all of the other reagents and primers).

