## Supplementary figures and images for "Removal of the catalytic subunit of DNA-protein kinase in the proximal tubules promotes DNA and tubular damage during kidney injury"

### Figure 1

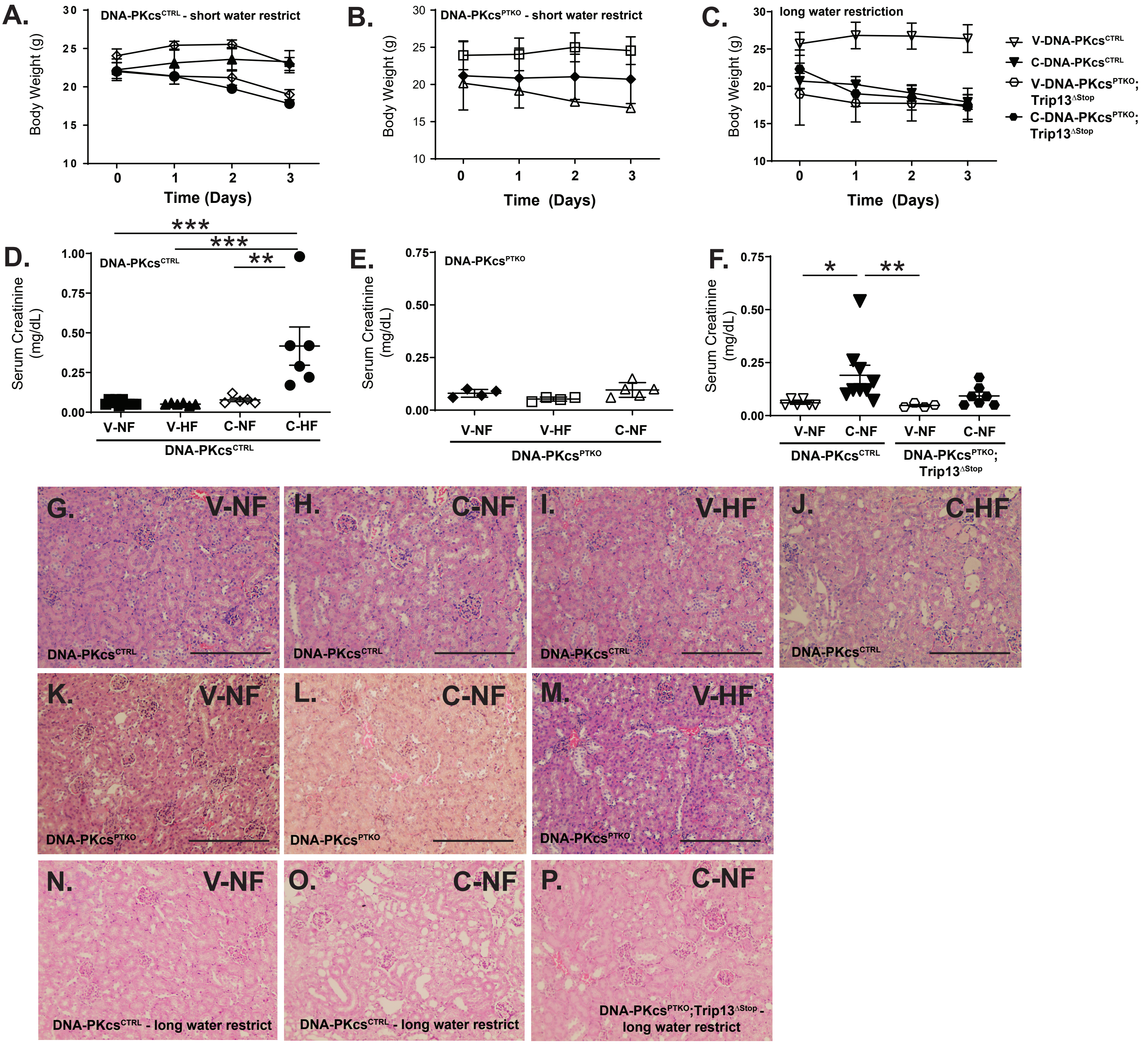

### Figure 2

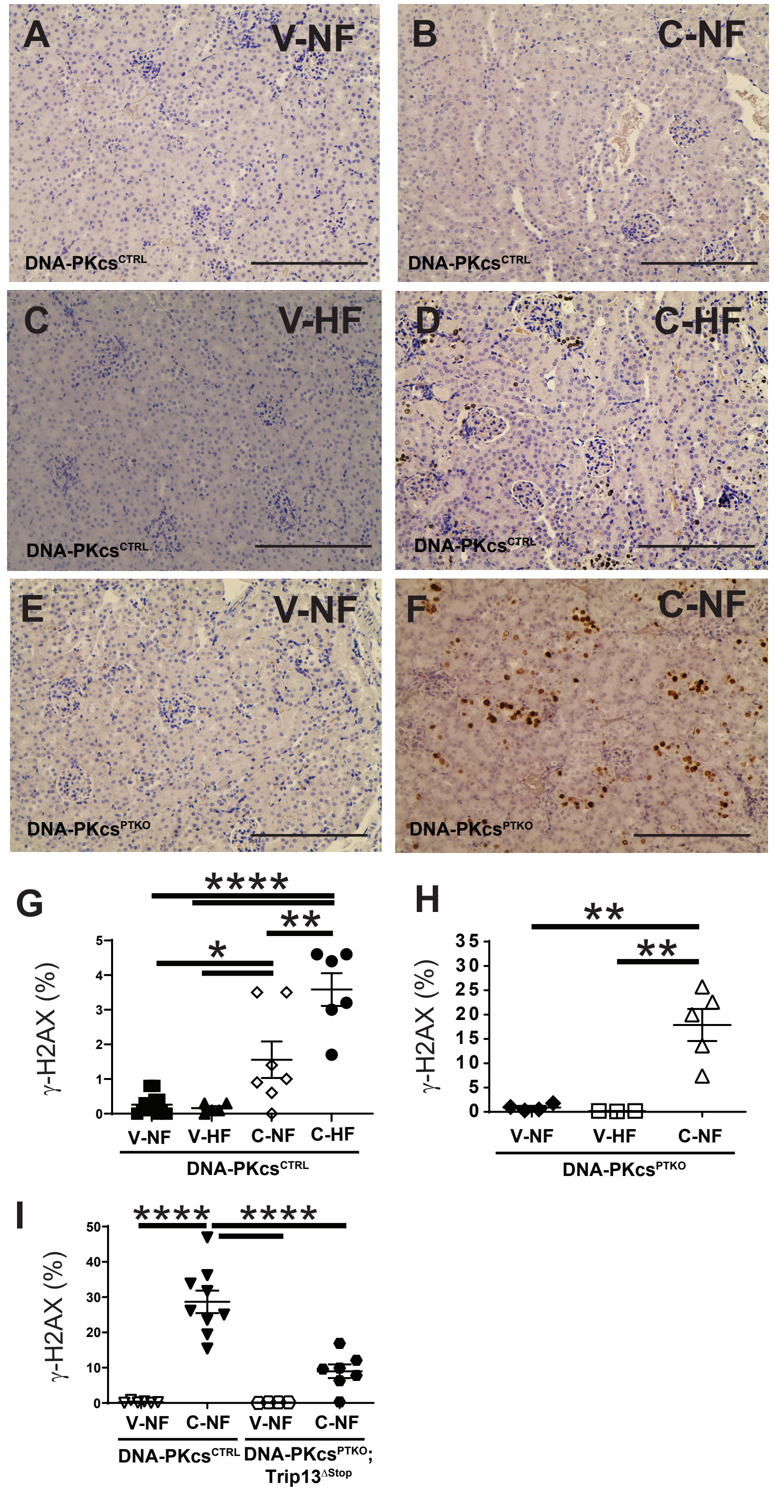

### Figure 5

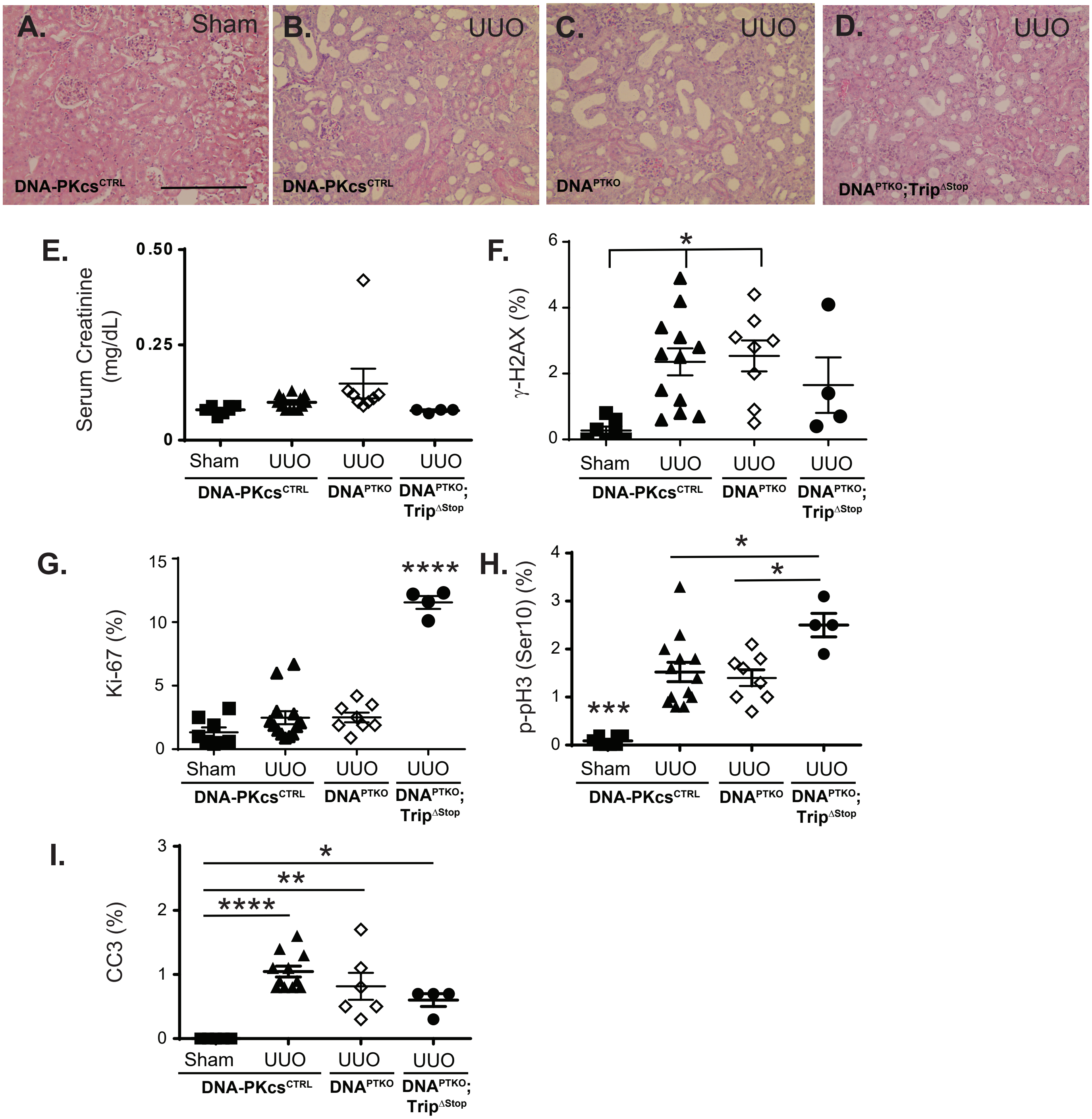
